# Feature Selection Enhances Peptide Binding Predictions for TCR-Specific Interactions

**DOI:** 10.1101/2024.10.11.617901

**Authors:** Hamid Teimouri, Zahra S. Ghoreyshi, Anatoly B. Kolomeisky, Jason T. George

## Abstract

T-cell receptors (TCRs) play a critical role in the immune response by recognizing specific ligand peptides presented by major histocompatibility complex (MHC) molecules. Accurate prediction of peptide binding to TCRs is essential for advancing immunotherapy, vaccine design, and understanding mechanisms of autoimmune disorders. This study presents a novel theoretical method that explores the impact of feature selection techniques on enhancing the predictive accuracy of peptide binding models tailored for specific TCRs. To evaluate the universality of our approach across different TCR systems, we utilized a dataset that includes peptide libraries tested against three distinct murine TCRs. A broad range of physicochemical properties, including amino acid composition, dipeptide composition, and tripeptide features, were integrated into the machine learning-based feature selection framework to identify key features contributing to binding affinity. Our analysis reveals that leveraging optimized feature subsets not only simplifies the model complexity but also enhances predictive performance, enabling more precise identification of TCR-peptide interactions. The results of our feature selection method are consistent with findings from hybrid approaches that utilize both sequence and structural data as input as well as experimental data. Our theoretical approach highlights the role of feature selection in peptide-TCR interactions, providing a powerful tool for uncovering the molecular mechanisms of the T-cell response and assisting in the design of more advanced targeted therapeutics.

## Introduction

The host adaptive immune response, primarily driven by the activation of T cells, orchestrates a precise and targeted defense by recognizing and responding to specific antigens (1, 2). T-cell receptors (TCRs) interact with peptide-major histocompatibility complex (MHC) through low-affinity, transient contacts, allowing them to identify the correct antigen while remaining sensitive to subtle molecular differences (3, 4). This low-affinity binding also allows for TCR cross-reactivity with diverse peptide sequences, broadening their recognition potential (2, 5). Accurately predicting peptide binding to specific TCRs is crucial for advancing immunotherapy and vaccine development, and for clarifying the underlying microscopic picture of immune response (5–8). However, this task remains very complex due to the immense variability of TCRs and peptides, in addition to the intricate nature of molecular mechanisms governing their binding affinities (9– 12).

There are multiple experimental techniques available for investigating TCR-peptide interactions, including crystallography (13), surface plasmon resonance (14), and yeast display systems (15–17). Compared to other methods, yeast display offers a unique high-throughput advantage, allowing the screening of a large number of peptide variants simultaneously, which enables the rapid identification of high-affinity interactions. In a recent study, a yeast surface display system was developed to screen highly diverse libraries of peptides presented by MHC molecules, identifying those capable of binding specific TCRs (5). By coupling this approach with deep sequencing, the sequence diversity of peptides recognized by different TCRs was mapped, which helped to uncover critical binding motifs and interactions. After multiple rounds of selection, their dataset was refined to include hundreds of high-affinity peptide-TCR interactions. This comprehensive experimental dataset opened opportunities for further computational techniques to clarify better the pertinent molecular features of TCR-peptide interactions.

The use of machine learning methods for predicting TCR-peptide interactions is a promising direction of studies that has the potential to overcome the limitations of traditional methods (18–23). By leveraging large datasets and incorporating structural, physicochemical, and sequence information, these models can learn the underlying principles that govern TCR specificity and binding affinity (24). Developing machine learning models to predict strong binder peptides for specific TCRs, however, involves several key challenges (24). A major one is the TCR cross-reactivity phenomenon when a single TCR can recognize and bind to multiple peptides (24–26). This property complicates the identification of true strong binders versus weaker ones, as a peptide that strongly binds to one TCR may bind weakly to another. One initial step for addressing this issue is to develop context-specific models to identify features that drive specificity in distinct functional scenarios, such as TCRs restricted to a common MHC allele that bind diverse peptide antigen sets. Additionally, quantification of such features in the context of the diversity to which strong binders are themselves identified, represents an important quantification of cross-reactivity within a given TCR system.

Recent computational frameworks (27–33), have made notable advancements in predicting TCR-peptide binding affinities. The Rapid Coarse-grained Epitope TCR (RACER) model is one particular example of a hybrid structural and sequence-based approach that uses a pairwise energy model trained on deep sequencing and crystallography data to identify strong and weak TCR-peptide binders. While RACER provides useful predictions, it relies on the availability of a relatively sparse set of experimentally determined TCR-peptide structures. Furthermore, RACER and other similar predictive models aim to benefit from leveraging biophysical features to subsequently require a reduced number of positive (1) and negative (0) examples in training. These models are trained on a collection of TCR-peptide systems and can handle variations in either the TCR or peptide. In our study, RACER was employed to analyze datasets restricted to the same MHC allele (all IE^k^), where it was previously used to resolve strong and weak TCR binding profiles. However, these models do not comprehensively evaluate all of the variability within a given binding class when, for example, we have a fixed TCR and a large number of confirmed strong and weak binding peptide sequences corresponding to that single TCR. Incorporating an ML-based classifier could complement models like RACER by extracting more nuanced, context-specific features that confer binding specificity. This synergistic approach could improve predictive accuracy within specific binding classes, enhancing our understanding of TCR-peptide interactions.

Our study aims to apply machine learning techniques with feature selection to improve the accuracy of TCR-peptide specificity prediction to identify motifs that most highly resolve strong and weak binders given available large-scale binding datasets. By identifying the most relevant features, among a comprehensive set of physicochemical features that determine binding interactions, our model effectively distinguishes between strong binders and weak binders. We apply various feature extraction techniques and examine the robustness of each approach. To test our theoretical method, the analysis is applied to three distinct peptide pools. The model’s ability to account for meaningful peptide variations that drive specificity is evaluated for each case, based on successful predictions of strong-and weak-binding TCR-peptide pairs.

## Materials and methods

### Dataset and data preprocessing

We employed a highly diverse set of peptide-MHC complexes derived from yeast-displayed peptide-MHC libraries, which includes three distinct types of murine TCRs: 2B4, 226, and 5cc7 (5). These TCRs were selected due to their distinct mechanisms of peptide recognition, which arise from variations in their structure, cross-reactivity, and interactions with MHC molecules (34). The peptide libraries for each TCR were subjected to multiple rounds of selection to enrich for TCR-binding peptides, which were subsequently analyzed using deep sequencing. The final dataset consisted of hundreds of unique peptide sequences, each characterized by specific TCR recognition motifs. In each dataset, each peptide sequence is characterized by a “Round 5” value, which refers to the fifth and final iteration in this selection process, where peptides that bind strongly to the TCR are identified. During each round of selection, the weaker binding peptides are gradually filtered out, and the frequency of peptides with stronger TCR affinities increases. Five rounds of affinity-based selection ultimately yield a list of sequences with their corresponding abundance, which indicates how many times that particular peptide was detected during sequencing and is proportional to their binding affinity.

Since the frequencies observed in Round 5 give a direct measure of how strongly and frequently each peptide binds to the TCR, we can classify peptides into strong binders (class 1) and weak binders (class 0) based on Round 5 selections by setting a threshold value calculated from the RACER model. To calculate the threshold value of Round 5 for separating strong binders from weak binders, a subset of 500 cases was selected from each dataset (5), with 140 instances designated as strong binders for training purposes. For each strong binder, a set of 1000 decoy sequences was generated by randomizing the peptide sequence and pairing it with the corresponding TCR structure as a comprehensive negative dataset that balanced the strong binders as was done previously (28), resulting in a total of 140, 000 decoy binders. The remaining cases were allocated for testing. We then applied RACER to compute thresholds that effectively separate the distributions of strong and weak binders for each TCR-pMHC case. The RACER model calculates binding energies by integrating high-throughput data from previously confirmed TCR-peptide interactions and crystal structures to train a residue-specific energy matrix. This energy matrix, combined with available structural templates, is used to quantify TCR-peptide binding affinities. For our peptides of interest, we utilized crystal structures with PDB IDs 3QIB, 3QIU, and 4P2R corresponding to 2B4, 226, and 5cc7 TCRs, respectively.

We predicted the binding energies and corresponding Z-scores for training and testing cases. To determine the threshold separating strong and weak binders, we analyzed histograms of the Z-scores for all 500 cases to identify the peaks of the distributions for both classes (Fig. 1). Fig. S1 presents the histograms for peptide libraries targeting 2B4, 226, and 5cc7 TCRs. The threshold was defined as the midpoint between the peaks of the strong and weak binder distributions. Finally, we ranked all 500 cases in the dataset based on their Z-score values which are obtained by the RACER model, and mapped these thresholds to the corresponding values reported in Round 5 of the dataset (5). The specific thresholds for each case are listed in Table 1. Using the calculated threshold values for each dataset, all peptides were partitioned into class 1 (strong binder) and class 0 (weak binder) cases. Since the dataset was highly imbalanced, with far more weak binders than strong binders, we performed controlled undersampling of the weak binders, which consisted of randomly selecting a subset of weak binders equal in size to the number of strong binders. This method helped mitigate the effects of class imbalance to improve the performance of the machine learning classifiers (35).

**Table 1.** Summary of selected peptide datasets associated with each TCR. Data obtained from Ref. (5). The thresholds for partitioning the data into strong (1) and weak (0) binders after five rounds of affinity-based selection were obtained using RACER model (27) (see text for details).

| TCR | Threshold | Peptides (0/1) |
| --- | --- | --- |
| 2B4 | 13 | 98/98 |
| 226 | 6 | 987/987 |
| 5cc7 | 23 | 234/234 |

**Fig. 1.**
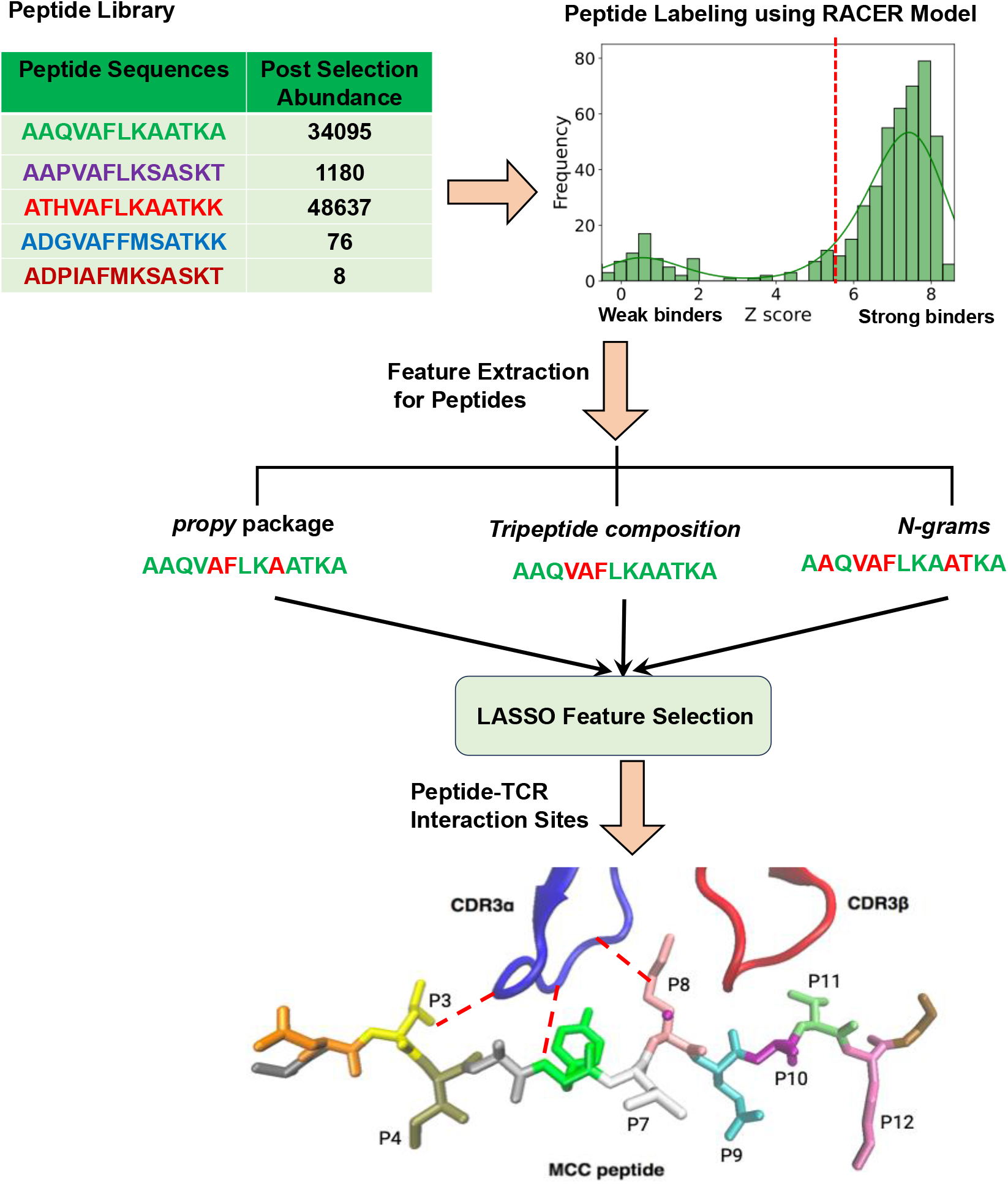
Schematic summary of the feature selection process for predicting peptide binding to T-cell receptors (TCR).

### Extraction of physicochemical features for peptides

To capture the physicochemical properties of peptides that are crucial for their interaction with TCRs, we extracted multiple features using primary amino acid sequence information. This is a critical step for implementing machine learning models. To evaluate the robustness of our feature selection technique, we employed three different methods for extracting physicochemical features from the sequence data.

#### propy package

First, we extracted a comprehensive set of physicochemical features from the amino acid sequence of each peptide using the *propy* package (36). These features are broadly categorized into different groups, including charge, amino acid composition, dipeptide composition, autocorrelations, chemical composition features, and sequence order information. The physicochemical features generated using *propy* package have been utilized in a wide range of machine learning models, including classification of antimicrobial peptides (37, 38) and predicting protein-protein interactions (39).

Among the features extracted by *propy*, amino acid composition and dipeptide composition are particularly important for understanding the interactions between TCRs and various peptides, as they provide insights into how specific residues and their combinations influence binding affinity (24). For a peptide of *L* residues, amino acid composition, which represents the fraction of each amino acid type, reads as

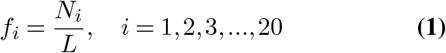

where *N*_*i*_ is the number of amino acids of type *i*. Since there are 20 amino acids, the amino acid composition comprises 20 features among the *propy* features.

Similarly, the dipeptide composition represents the fraction of each possible dipeptide within the peptide, calculated as:

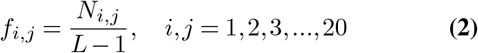

where *N*_*i*,*j*_ is the number of dipeptides consisting of amino acids of type *i*,*j*. Consequently, the dipeptide composition contributes 20^2^ = 400 distinct features to the set of *propy* features.

#### Tripeptide Composition

Tripeptide composition represents the fraction of each possible tripeptide (formed by three consecutive amino acids) within a peptide sequence. The tripeptide composition is calculated as:

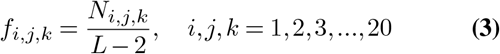

where *N*_*i*,*j*,*k*_ is the number of tripeptides containing amino acids of type *i*,*j*,*k*. Tripeptide composition, which comprises 20^3^ = 8000 features, provides deeper insight into the peptide’s structure by capturing the frequency of every unique combination of three consecutive amino acids. Since the *propy* package does not provide tripeptide composition features by default, we extracted these features separately.

#### N-gram language model

A sequence of amino acids, whether forming a short peptide or a large protein, can be viewed as a text document, where amino acids function as the fundamental units, analogous to words (40). Text mining and natural language processing have been previously employed for bioinformatics applications such as protein clustering and classification, protein-protein interaction (PPI) prediction, protein folding analysis, and non-coding RNA identification (41, 42).

To analyze amino acid sequences using natural language processing methods, we can use the *N* -gram language model, which is a probabilistic model used to predict the next item in a sequence based on the preceding items. A *N* -gram is a contiguous sequence of *N* items from a given sequence of text. In our context, each amino acid represents an item (analogous to a word), and a *N* -gram would be a sequence of *N* amino acids. While *propy* can efficiently compute the frequency of single amino acids and dipeptides, the resulting dipeptide frequencies tend to be sparse, as many dipeptides may not appear in a given peptide. By incorporating common *N* -grams including unigrams (single amino acids), bigrams (pairs of amino acids), and trigrams (triplets), the model goes beyond mere composition analysis and captures the sequential order and local motifs within peptides (43). Moreover, the robustness of the overall predictive model can be enhanced by combining different types of amino acid composition, including unigrams, bigrams, and trigrams. This approach ensures that if one feature set fails to capture critical patterns, the other can compensate, leading to a more comprehensive and accurate analysis of peptide-TCR interactions.

Since all peptides are composed of 20 standard amino acids, the maximum vocabulary sizes for unigrams, bigrams, and trigrams are 20, 20^2^ = 400, and 20^3^ = 8000, respectively. This creates a fixed-size vocabulary that can be represented as a numerical feature vector, where each element corresponds to the frequency or presence of a specific *N* -gram in the sequence. The process of vectorizing a peptide sequence using the *N* -gram approach begins by breaking down each peptide into unigrams, bigrams, and trigrams, which serve as the fundamental building blocks of the sequence. Next, a complete vocabulary is composed of all possible *N* -grams that can occur within the sequence. Once the vocabulary is established, the sequence is vectorized by converting the frequency of each *N* -gram into a numeric vector. The resulting vectorized transformation, which has lower sparsity compared to features generated by *propy*, enables efficient processing of peptide sequences by machine learning algorithms.

### Data Normalization

For each peptide, the quantitative values of the physicochemical properties extracted from the methods described above have different numerical scales. It is important to initially re-scale all these values to fall between 0 and 1 so that every property is considered with a similar weight. To normalize this quantity to be in the range 0 and 1, we use the following re-scaling expression,

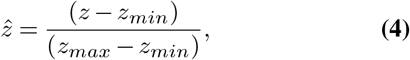

where *z* is the original value of the physicochemical property, *z*_*min*_ and *z*_*max*_ are limiting values for this property for all considered proteins, and 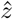 is the normalized one that is specifically utilized in the analysis. It is important to note that to prevent leakage from the training set to the test set, we performed data normalization only after splitting the datasets into training and test sets.

### Feature Selection

In studying TCR-peptide interactions, our primary goal is to identify which specific physicochemical features – such as amino acid properties or sequence motifs – are most important for distinguishing between strong and weak binders. However, the extracted feature set often consists of high-dimensional data, meaning the number of features may exceed the available data, with some being irrelevant or highly correlated. Using all these features with-out selection can result in overfitting, where the model learns noise rather than meaningful correlations, reducing its predictive performance (37). To mitigate this issue, we employ LASSO (The Least Absolute Shrinkage and Selection Operator) techniques that mathematically assign zero weights to irrelevant or redundant features (37, 44). The overview of our feature selection procedure is presented in Fig. 1.

## Results

The relative significance of various physicochemical features distinguishing strong binders from weak binders in 2B4, 226, and 5cc7 peptide libraries are presented in Figs. 2, 3, and 4, respectively. We performed feature selection using three categories of properties: *propy* features, tripeptide composition, and N-gram language model (unigram, bigram, and trigram). This multi-faceted feature generation approach enabled us to extract key patterns and properties that significantly influence TCR binding behavior.

**Fig. 2.**
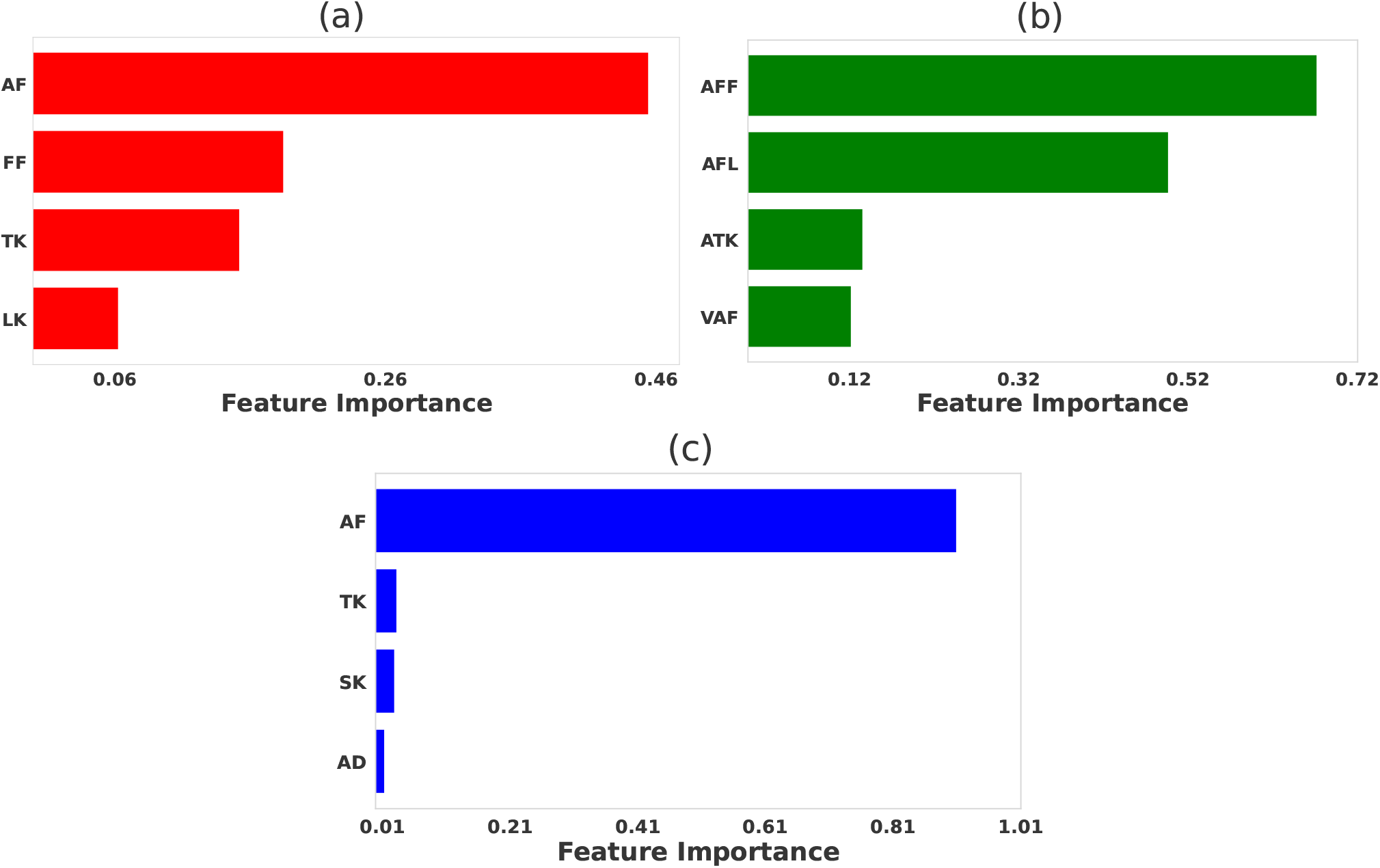
Comparative significance of various physicochemical features in differentiating strong and weak peptide binders of the 2B4 TCR. LASSO feature selection was performed using features generated using (a) *propy* (including dipeptide composition), (b) tripeptides composition, (c) N-gram language model (incorporating unigram, bigram, and trigram). In each case, the LASSO hyperparameters were set to be *λ* = 0.005.

**Fig. 3.**
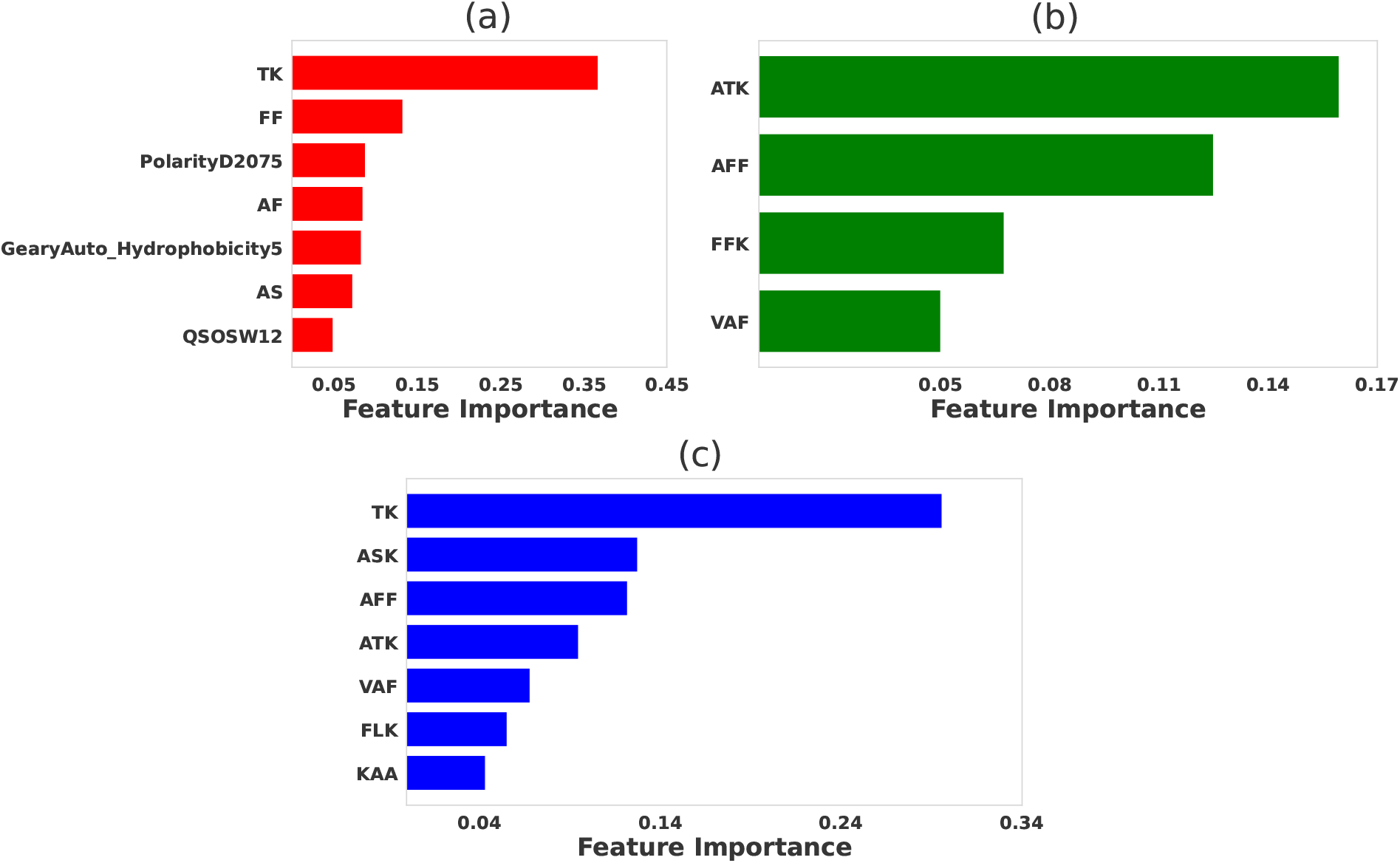
Comparative significance of various physicochemical features in differentiating strong binder and weak binder for peptides targeting 226 TCR. LASSO feature selection was performed using features generated using (a) *propy* (including dipeptide composition), (b) tripeptides composition, (c) N-gram language model (incorporating unigram, bigram, and trigram). For LASSO the hyperparameters were set to be *λ* = 0.015, *λ* = 0.015, and *λ* = 0.01, respectively.

**Fig. 4.**
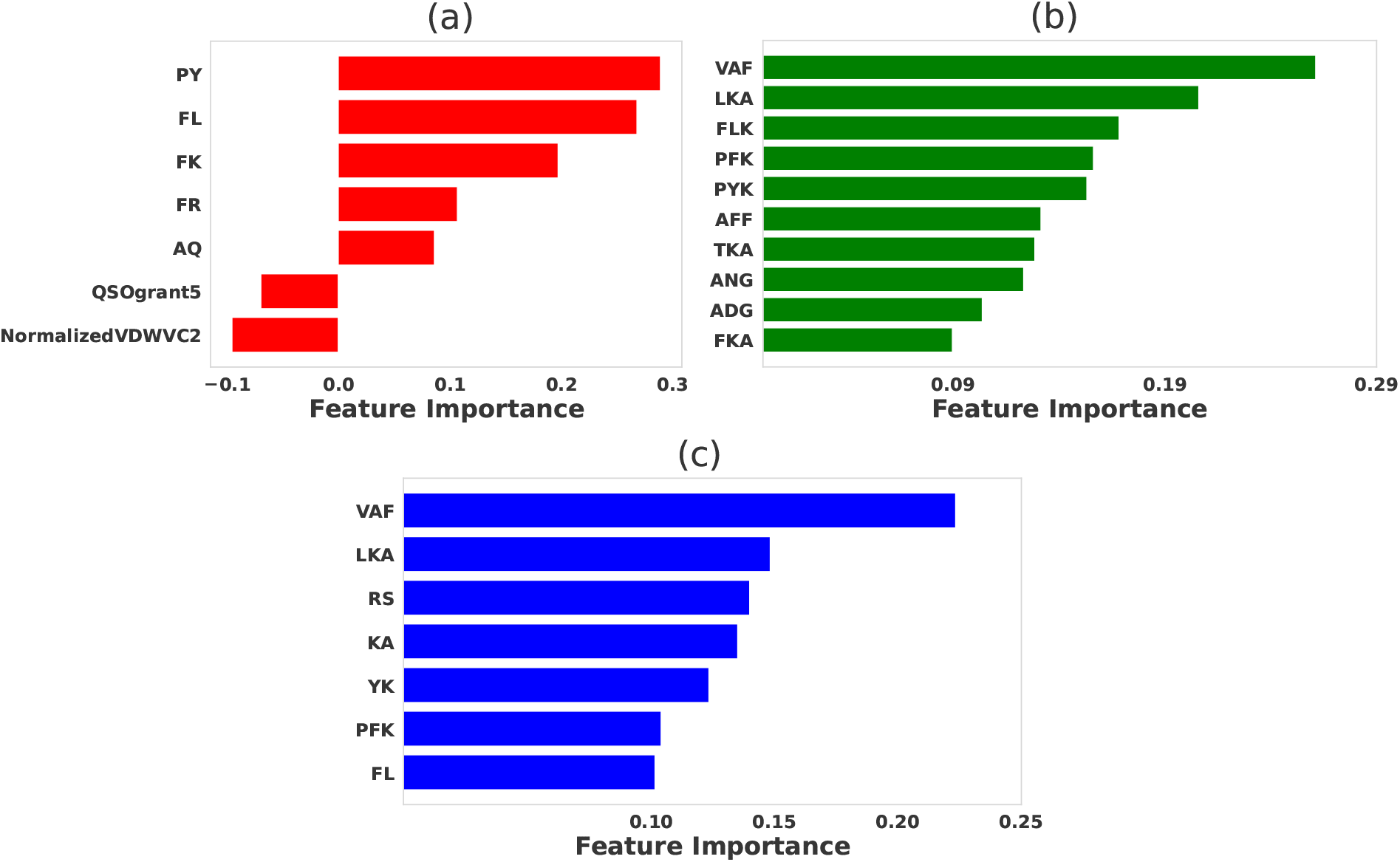
Comparative significance of various physicochemical features in differentiating strong binder and weak binder for 5cc7 peptides. LASSO feature selection was performed using features generated using (a) *propy* (including dipeptide composition), (b) tripeptides composition, (c) N-gram language model (incorporating unigram, bigram, and trigram). For LASSO the hyperparameters were set to be *λ* = 0.01 for all cases.

### Features Selection for 2B4 data

Our feature selection method for 2B4 data yields different but in many aspects overlapping selected features that contribute to strong binder peptides. Among features generated from the *propy* tool, the most important dipeptide compositions such as ‘AF’, ‘FF’, ‘TK’, and ‘LK’ likely represent amino acid pairs that significantly enhance peptide stability or affinity to the TCR [see Fig. 2 (a)]. Similar motifs are predicted when tripeptide compositions are utilized in the feature selection method, as shown in Fig. 2 (b). Specifically, the tripeptide motif ‘AFF’ can be broken down into two dipeptides ‘AF’ and ‘FF’, both of which are captured by the *propy* method. The N-gram method also yields similar results, although selected features do not fully overlap with the results of other methods [see Fig. 2 (b)].

This observation is strongly supported by the experimental data, which highlights the amino acid preferences at key TCR contact positions (P3, P5, and P8) during peptide-MHC interactions (5). Notably, positions like P3 show a clear preference for aromatic residues such as phenylalanine (F) and tyrosine (Y), aligning with the dipeptides ‘AF’ and ‘FF’, and the tripeptide ‘AFF’, identified in our study. The overlap between dipeptides ‘AF’, ‘FF’ and tripeptides ‘AFF’, ‘AFL’ across different feature selection methods demonstrates that these motifs are important for strong binders. This consistency is also reflected in the N-gram results, where motifs such as ‘AF’ dominate, highlighting the key structural patterns that underlie strong TCR-peptide interactions.

### Features Selection for 226 data

For the 226 TCR dataset, our feature selection method also identified several key physicochemical features that distinguish strong binders from weak binders, as presented in Fig. 3. Among the features generated by the *propy* tool [see Fig. 3(a)], dipeptide compositions such as ‘TK’, ‘FF’, ‘AF’ emerged as the most important contributors to peptide-TCR affinity. These motifs likely play critical roles in stabilizing peptide-TCR interactions by complementing specific amino acid residues on the TCR. Structural analysis of the 226 TCR-pMHC (PDB ID:3QIU) reveals that the ‘TK’ motif contributes to electrostatic interactions, and ‘FF’ and ‘AF’ reinforce hydrophobic interactions that enhance the stability of the peptide-MHC complex. Definitions of other selected *propy* features, including PolarityD2075, GearyAuto_Hydrophobicity5, and QSOSW12 are presented in Table S1 in the supplementary information.

Specifically, in the ‘TK’ motif, the weakly acidic threonine residue (‘T’ at P8) can interact via hydrogen bonding with asparagine on the TCR’s CDR3*β* loop. This interaction is identified through the contact map generated from the 3QIU crystal structure with a maximum distance (*r*_max_ = 8.5 Å) (Fig. S5(c),(d)). Moreover, for the ‘FF’ and ‘AF’ motifs, which do not exhibit these features in the original peptide, the RACER-derived pairwise amino acid energy matrix (Fig. S4(b)) predicts a favorable interaction between phenylalanine (‘F’) and alanine (‘A’). Similarly, alanine is predicted to favorably interact with proline (‘P’), methionine (‘M’), and phenylalanine (‘F’). These findings suggest that the ‘F’ residues in the peptide engage in favorable interactions with ‘A’ and other hydrophobic residues on the TCR. These hydrophobic interactions contribute to the stability of the peptide-TCR complex, underscoring the importance of the ‘FF’ and ‘AF’ motifs in facilitating binding through hydrophobic contacts.

Tripeptides such as ‘ATK’, ‘AFF’, and ‘FFK’ were identified as highly significant for distinguishing strong binders (see Fig. 3(b)). These tripeptides can be deconstructed into dipeptides like ‘TK’, ‘FF’, and ‘AF’, which contain residues that are also prominent in the dipeptide analysis. The similarity between the tripeptide and dipeptide results reinforces the importance of these specific motifs, suggesting that key residues such as lysine (‘K’) and phenylalanine (‘F’) play central roles in facilitating hydrogen bonding and electrostatic interactions with the TCR’s complementarity-determining regions (CDRs). For example, lysine (‘K’) in the ‘ATK’ and ‘FFK’ motifs likely contributes to salt bridge formation, enhancing electrostatic interactions between the peptide and TCR. Similarly, phenylalanine (‘F’) in the ‘AFF’ and ‘FFK’ motifs strengthens binding through hydrophobic interactions, which help stabilize the peptide-TCR complex within the MHC groove.

Furthermore, feature selection based on the N-gram language model [see Fig. 3(c)] revealed a strong overlap with the amino acid patterns identified in both the dipeptide and tripeptide analyses. The most important features included ‘TK’, ‘ASK’, ‘AFF’, and ‘ATK’, which closely correspond to amino acids found at critical TCR contact points. This consistency across different feature selection methods reinforces the importance of these motifs in contributing to strong peptide-TCR interactions. The recurrence of ‘TK’, in particular, highlights the role of lysine in driving strong electrostatic interactions, while ‘AFF’ and ‘FFK’ emphasize the contribution of hydrophobic residues like phenylalanine (F) in maintaining binding affinity and structural stability.

### Features Selection for 5cc7 data

Our feature selection approach, trained using features generated from *propy* tool, predicts that ‘PY’, ‘FL’, ‘FK’, and ‘FR’, are crucial in determining peptide stability and TCR binding affinity [see Fig. 4(a)]. These dipeptides are likely involved in stabilizing peptide-MHC interactions, with ‘FL’ and ‘FK’ contributing to hydrophobic and polar interactions, respectively. Hydrophobic residues such as phenylalanine (F) are known to form important nonpolar contacts that help to stabilize the peptide in the TCR binding groove, enhancing binding affinity. Definitions of the acronyms for other selected *propy* features — QSOgrant5 and NormalizedVDWVC2 — are presented in Table S1 of the supplementary information.

When the tripeptide compositions are considered [see Fig. 4(b)], motifs like ‘VAF’, ‘LKA’, and ‘FLK’ emerged as highly significant. These tripeptides suggest a combination of hydrophobic, polar, and charged interactions, which enhance the binding potential by promoting stable contacts of different natures with the TCR. For instance, ‘LKA’ features a combination of leucine (‘L’) and alanine (‘A’), hydrophobic residues, and lysine (‘K’), a positively charged residue, both of which are known to interact favorably with the TCR’s (PDB ID: 4P2R) binding pocket through hydrophobic and electrostatic interactions. Notably, the ‘LKA’ motif does not appear in the original peptide contact map generated with a maximum distance (*r*_max_ = 8.5 Å) (Fig. S5(e),(f)). However, analysis of the energy matrix (Fig. S4(c)) reveals that leucine (‘L’) has high affinity for proline (‘P’), alanine (‘A’), and phenylalanine (‘F’), suggesting potential hydrophobic interactions with these residues on the TCR. Similarly, lysine (‘K) shows high affinity for tryptophan (‘W’), alanine (‘A’), phenylalanine (‘F’), and leucine (‘L’), indicating possible favorable interactions with these residues. Furthermore, alanine (‘A’), due to its small side chain, provides structural flexibility, allowing optimal positioning of neighboring residues for interaction. These findings suggest that the ‘LKA’ motif may enhance peptide-TCR interactions through hydrophobic and electrostatic interactions, as indicated by the energy matrix analysis, even though these interactions are not apparent in the contact map.

The N-gram language model (Fig. 4(c)) further enlightened the importance of these motifs by identifying similar patterns. Features like ‘VAF’, ‘LKA’, ‘RS’, and ‘KA’ were among the most important for distinguishing strong binders, reflecting the same key interactions seen with the dipeptide and tripeptide compositions. The prevalence of hydrophobic residues such as valine (V), phenylalanine (F), and leucine (L) in these motifs emphasize the critical role in stabilizing the peptide-MHC-TCR complex.

### Prediction of strong vs weak binders using selected features

After selecting the essential physicochemical features from each peptide dataset, we aim to leverage these attributes to accurately and reliably predict strong and weak binders for each TCR type through logistic regression models. Table 2 summarizes key performance metrics, including Accuracy, Recall, F1 Score, Matthews Correlation Coefficient (MCC), and AUC (Area Under the ROC Curve), averaged over 10 cross-validation sets with an 80/20 train/test split for each fold.

**Table 2.** Performance comparison of feature selection methods for three TCR data sets. Metrics include Accuracy, Recall, Matthews Correlation Coefficient (MCC), F1 Score, and AUC for trained baseline models (Logistic Regression) using selected features from LASSO. Values reflect the average across 10 cross-validation sets, with an 80/20 train/test split for each fold.

| TCR Data | Feature Category | Accuracy | Recall | F1 Score | MCC | AUC |
| --- | --- | --- | --- | --- | --- | --- |
| 2B4 | propy | 0.94 | 0.94 | 0.93 | 0.87 | 0.96 |
|  | tripeptide | 0.96 | 0.96 | 0.96 | 0.92 | 0.97 |
|  | N-gram | 0.96 | 0.96 | 0.96 | 0.93 | 0.98 |
| 226 | propy | 0.77 | 0.77 | 0.79 | 0.55 | 0.78 |
|  | tripeptide | 0.74 | 0.74 | 0.76 | 0.49 | 0.66 |
|  | N-gram | 0.73 | 0.73 | 0.76 | 0.48 | 0.7 |
| 5cc7 | propy | 0.82 | 0.82 | 0.83 | 0.65 | 0.83 |
|  | tripeptide | 0.85 | 0.85 | 0.86 | 0.71 | 0.87 |
|  | N-gram | 0.85 | 0.85 | 0.85 | 0.79 | 0.84 |

For the 2B4 dataset, all three selected feature categories (*propy*, tripeptide composition, and N-gram) performed exceptionally well, with predictive accuracy ranging from 0.94 to 0.96 and an AUC reaching up to 0.98. These findings indicate that for this dataset, the selected features were highly informative, resulting in predictive models that perform well in identifying strong binders with high precision. The strong performance of the models for the 2B4 dataset can be attributed to the lower cross-reactivity of 2B4 i.e. it binds to a narrower range of peptides compared to more flexible TCRs, making the binding interactions easier to model and predict. Furthermore, a smaller dataset (98 peptides) with clear sequence patterns provides the machine-learning models with less variability to account for, resulting in higher accuracy and AUC values.

In contrast, the 226 datasets demonstrated somewhat lower overall performance across all feature methods. Accuracy and AUC values were notably lower, with *propy* yielding the highest performance at 0.77 accuracy and 0.78 AUC, while the tripeptide and N-gram methods scored marginally lower. The relatively low MCC values (0.55 for *propy* and below 0.50 for others) suggest that the model’s predictions are less consistent for this dataset. This result is likely due to the existence of more complex or less distinguishable features between strong and weak binders. The relatively lower performance of the models for the 226 dataset can be attributed to several factors related to the biological properties of the 226 TCR and the complexity of its dataset. The 226 TCR is known for its high degree of cross-reactivity (34), meaning it can recognize and bind to a much wider range of peptide sequences than more specific TCRs like 2B4. This broad recognition profile introduces greater variability in the peptide sequences classified as binders and non-binders, making it harder for machine learning models to identify clear patterns that distinguish strong from weak binders. Thus, the larger size of the 226 dataset, which includes 987 peptides, increases the diversity of peptide sequences.

For the 5cc7 dataset, however, performance is intermediate, with accuracy values ranging from 0.82 to 0.85 and an AUC as high as 0.87 for the tripeptide method. Here, the MCC values indicate that the models were relatively effective, with the N-gram method achieving the highest MCC (0.79), suggesting that it provided a more balanced prediction between strong and weak binders compared to the other methods. The F1 scores consistently reflect solid performance in identifying true strong binders, particularly with the tripeptide method (*F*_1_ = 0.86). The moderate performance of the models for the 5cc7 dataset can be explained by the balance between specificity and cross-reactivity in the 5cc7 TCR and the size of the dataset. Unlike the highly specific 2B4 TCR or the highly cross-reactive 226 TCR, 5cc7 exhibits an intermediate level of specificity. It binds to a moderately diverse set of peptides, leading to less sequence variability than 226 but more than 2B4.

It is important to highlight that while our datasets include an equal number of strong and weak binders, the overall peptide data are highly imbalanced in favor of weak binders over strong ones. Quantitatively, for a peptide of *L* residues, the total number of possible peptide sequences are 20^*L*^, and an overwhelming majority of these sequences are weak binders. This discrepancy presents significant challenges in accurately predicting peptide specificity. However, despite these challenges, the close alignment between our *F* 1-score and recall metrics (Table 2) indicates that the model achieves balanced performance in handling false positives (FP) and false negatives (FN). The balance between Recall and *F* 1-score is especially critical in this context, where accurately identifying strong binders is essential, but misclassifying weak binders as strong could lead to a false sense of antigen coverage for a particular therapy. The fact that both metrics are comparable across datasets and feature selection methods indicates that the models are balanced in their sensitivity and specificity and are robustly selecting relevant features to resolve strong and weak binders (39).

### Comparison with the RACER model

Our feature selection-based approach for predicting TCR-peptide binding, like several before it (30, 31, 33, 45–47), is purely sequence-based, relying on the selection of key features derived from amino acid sequences. By focusing on the sequence characteristics of peptides, we identified key dipeptide and tripeptide motifs that are enriched in strong binders. These selected motifs, without relying on detailed structural information, were essential for distinguishing between high- and low-affinity peptide antigens for specific TCR. In contrast, the RACER model adopts a hybrid approach by combining sequence data with structural insights to predict TCR-peptide binding affinities (21, 27, 28, 48, 49). RACER utilizes a pairwise energy framework, integrating residue-specific energy matrices derived from highthroughput data on experimentally confirmed TCR-peptide interactions, along with crystal structures of these complexes. The structural templates provided by crystal data allow RACER to quantify the binding energy with greater precision by modeling the physical interactions at atomic resolution. After identifying motifs enriched in strong binders, we then aimed to apply RACER’s pairwise energy framework to test our sequence-based approach. This allowed us to pinpoint the specific positions within the peptide sequence where these motifs are predicted to have the most significant impact on binding energy.

To determine the positions of the selected features, we performed *in silico* mutation in all two-adjacent (selected dipeptide motifs) and three-adjacent (selected tripeptide motifs) amino acids at every peptide amino acid position containing the selected features. We then used RACER to estimate the binding energy for each mutant peptide. The binding energies for all possible mutant peptides are plotted for selected dipeptides (Fig. S2) and tripeptides (Fig. S3). We then compared the binding energy of each oligopeptide motif located at each position to the baseline binding energy of the wild-type (WT) TCR-peptide (WT given by the black dashed line in Figs. S2 and S3). If a mutated TCR-peptide showed increased binding energy (above the dashed line), it indicates that the mutation enhanced the binding affinity above that of the WT (strong) binding peptide, suggesting that the underlying importance of the selected dipeptide at that specific position. On the other hand, mutation may also result in significantly lower predicted affinities.This decrease indicates that certain substitutions disrupt key interactions necessary for strong binding, effectively identifying sequences that function as weak binders. By recognizing these sequences, we not only validate the specificity of our selected motifs but also enhance our understanding of the structural and sequence determinants that diminish binding affinity. This dual identification of both strong and weak binders underscores the robustness of our approach in mapping the landscape of TCR-peptide interactions.

For example, Fig. S2(a) shows that for 2B4, dipeptide ‘AF’ at positions (2,3), (3,4), and (5,6) increased binding affinity, with a particularly significant increase at positions (5,6). Although all three TCRs retain a WT-like TCR recognition motif, each TCR exhibits some variation in positional preference (Fig. 5). For instance, whereas 2B4 can recognize Lysine at position P8 (P5 in Ref. (5)), 5cc7 accommodates Leucine and Arginine at P8. The P6 (P3 in Ref. (5)) TCR contact position showed the least variance across all three TCRs, with either Phenylalanine or Valine being required for 2B4 and 5cc7, and Phenylalanine, Lysine, or Arginine being required for 226. As previously reported (5), 226 demonstrated a greater degree of cross-reactivity, being able to recognize 897 unique peptide sequences. The larger number of peptides recognized was largely due to a higher tolerance for substitutions at TCR-neutral and MHC-contacting residues, such as position P9 (Fig. 5(b)).

**Fig. 5.**
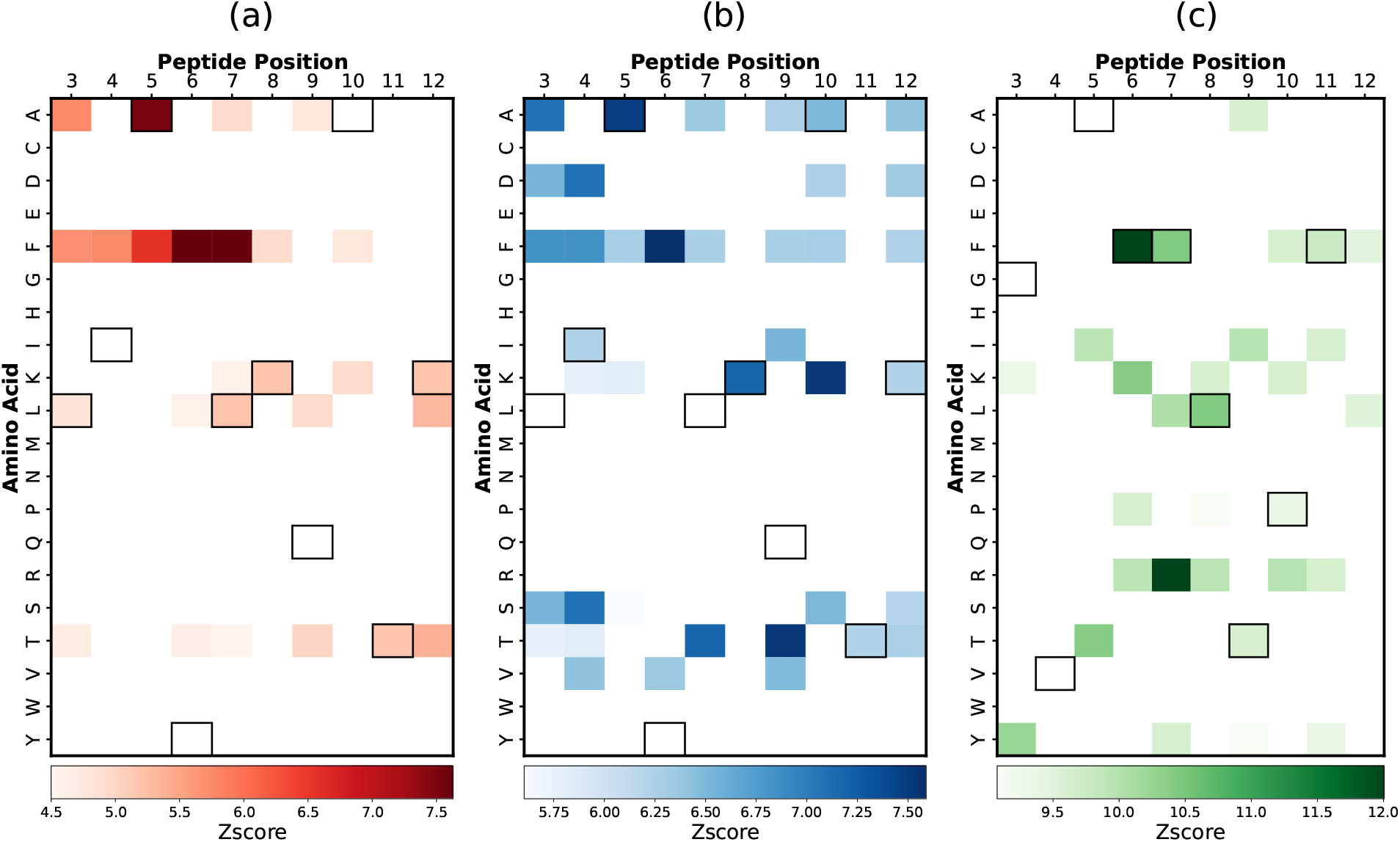
Predicted heatmaps using RACER model for (a) 2B4, (b) 226, and (c) 5cc7 peptide libraries. The sequence for the peptide is represented via outlined boxes.

Combining the predictions identified in our feature selection framework with RACER-predicted position-specific information provides an opportunity to construct heatmaps (Fig. 5) enriched in beneficial dipeptide compositions that maximally resolve strong and weak binding peptides. These results can be directly compared to those from the original work by Birnbaum et al. (5), which provided a similar description of binding motifs acquired experimentally. Notably, while their work identified *single amino acid* hotspots indicative of strong binders by analyzing the abundance of amino acids in strong-binding peptides, our approach focuses on *dipeptide motifs*, identifying them based on their enrichment in strong binders relative to non-binders.

This methodological difference is evident when considering anchor residues like P4 and P12, which are restricted to isoleucine, leucine, valine, and lysine, respectively. While lysine is ubiquitous among all strong binders, it is similarly present in weak binders and thus does not emerge as a distinguishing amino acid at P12 in our analysis (Fig. 5). This underscores one way in which conserved residues might mask a model’s discriminatory power were it to identify those features as important for strong binding. In addition to this, in the experimental data from (50), heatmaps are generated after three round of selection, whereas in our approach, we generate heatmaps after five round of selection.

Despite these challenges, the key motifs identified through our sequence-based feature selection are corroborated by RACER’s binding energy predictions. Particularly in the case of the 2B4 TCR, where we predict the enrichment of specific motifs such as the ‘AFF’ motif. RACER’s energy calculations confirm that these motifs contribute significantly to binding affinity (as indicated by larger interaction values with complementary amino acids ‘WSQ’ in the 2B4 TCR CDR3*β* domain, and ‘RA’ and ‘G’ in the CDR3*α* domain), aligning with experimental observations. The 2B4 TCR seems well-characterized by one important motif, as evidenced by the peptide position curves showing a single sharp peak corresponding to a small number of features (Fig. S2(a) and S3(a)). In contrast, the 226 and 5cc7 TCRs display different binding characteristics, highlighting the unique specificity of each TCR. For 5cc7, we observe lower intensity compared to 2B4, with a wide and smooth peak across a larger number of features (Fig. S2(c) and S3(c)). For 226, we observe sharp peaks at different positions for a large number of features, which aligns well with the high cross-reactivity we previously mentioned for 226 (Fig. S2(b) and S3(b)). In both 2B4 and 226, we observe the importance of features containing phenylalanine (‘F’) in the first part of the peptide across positions P3 to P7. This is because the peaks for dipeptide and tripeptide motifs containing ‘F’ are prominent in these positions. When considering dipeptide and tripeptide motifs, ‘F’ appears in all positions P3–P7, whereas in single amino acid analysis, ‘F’ does not appear in all positions since only the best location is selected. This indicates that ‘F’ can be a very important feature when combined with other amino acids in motifs. These findings underscore the power of our model in identifying critical dipeptide and tripeptide motifs, which are more informative than single amino acid motifs, thereby enhancing predictive performance and providing deeper insights into TCR-peptide interactions.

## Discussion

The interaction between T-cell receptors (TCRs) and peptide-MHC complexes is a critical component of the adaptive immune system, enabling T cells to detect and respond to specific antigens. This process, however, is complicated by the TCR cross-reactivity, where a single TCR can recognize multiple peptide sequences. Understanding cross-reactivity is important as many TCRs are known to confer coverage across many distinct peptide systems, but it also presents a major challenge in predicting specific TCR-peptide binding affinities, which is a critical need in immunotherapy and vaccine design. In this study, we aim to leverage machine learning techniques with refined feature selection to improve the accuracy and generalizability of TCR-peptide interaction predictions. Our findings show that focusing on specific physicochemical features significantly enhances the model’s ability to distinguish between strong and weak binders, offering new insights into the molecular mechanism of TCR recognition. The number of peptides in our final dataset reflects the varying specificity and cross-reactivity of the TCRs, which in turn explains the differences in model performance. The 2B4 dataset, with only 98 peptides, highlights the high specificity of the 2B4 TCR, leading to clearer binding patterns and superior model performance. In contrast, the 226 dataset, which includes 987 peptides, highlights the TCR’s greater cross-reactivity, making binding patterns more complex and harder to capture with our feature selection method, resulting in lower performance metrics. The 5cc7 dataset, with 234 peptides, demonstrates intermediate specificity and moderate cross-reactivity, aligning with its intermediate model performance. This variation in dataset sizes reflects the inherent biological properties of each TCR, with more specific TCRs resulting in smaller datasets and higher model performance. We employed the LASSO feature selection method to extract meaningful physicochemical properties from the peptide sequences to identify key features that contribute to binding affinity, including amino acid composition, dipeptide frequency, and tripeptide motifs. Among these, dipeptide compositions and tripeptide compositions emerged as particularly important, consistently ranking among the most predictive for distinguishing strong from weak binders across the different TCR datasets. This suggests that the arrangement of amino acids in short peptide sequences plays a crucial role in TCR recognition. This optimized feature set provides a robust foundation for understanding peptide-TCR interactions and highlights the importance of tripeptides in the subsequent analysis.

The importance of selected tripeptides in TCR-peptide binding can be understood by examining the molecular interactions between peptide residues and the CDR3*α* and CDR3*β* loops of the TCR, as illustrated in Fig 1. It is known that a single amino acid in the peptide can simultaneously interact with residues from both the CDR3*α* and CDR3*β* regions (34). For example, if we consider a symbolic tripeptide sequence like ‘LTP’ the first residue, ‘L’, may form contacts with both CDR3*α* and CDR3*β*, providing a dual interaction site. In contrast, the second and third residues, ‘T’ and ‘P’, may predominantly interact with only CDR3*β*. This picture highlights how specific residues within a tripeptide can influence the binding strength by creating multiple interaction points, making tripeptides like ‘LTP’ particularly important for determining binding affinity. The ability of certain tripeptides to establish multiple points of contact contributes to the overall specificity and affinity of TCR recognition.

While our sequence-based approach successfully identifies key dipeptide and tripeptide motifs enriched in strong binders, it has certain limitations. Our purely sequence-driven model may miss rare or unconventional motifs and struggle in cases of extreme TCR cross-reactivity. Additionally, our findings are derived from a relatively limited set of TCR-peptide interactions, which may limit the generalizability of the identified motifs across all TCRs, particularly those with unique binding preferences. Moreover, certain TCRs may prioritize interactions with MHC residues over peptides, a factor that our current model does not fully address. To overcome these limitations, future work will explore hybrid models that integrate structural insights, allowing for more accurate predictions of TCR-peptide dynamics. Despite these challenges, this approach is able to extract meaningful motifs for resolving TCR specificity based on TCR and peptide primary sequences. Future work will be directed at using these learned features to train a classification model for identifying strong binding pairs from a variety of possible TCR and peptide test sequences.

## Author Contributions

HT, ZSG, ABK, and JTG designed the research. HT and ZSG performed the research. HT, ZSG, ABK, and JTG analyzed the data and wrote the paper. All authors approve of the final version.

## Data Availability Statement

The datasets analyzed for this study can be found in the RACER github repository https://github.com/XingchengLin/RACER/tree/main/raw_data.

The code used in this study can be found in https://github.com/TAMUGeorgeGroup/Feature_Selection_TCR-Specific_Interaction.git

## References

1. Janeway Ca. The immune system in health and disease. http://www.garlandscience.com, 2001.

2. Michael E Birnbaum, Shen Dong, and K Christopher Garcia. Diversity-oriented approaches for interrogating t-cell receptor repertoire, ligand recognition, and function. Immunological reviews, 250(1):82–101, 2012.

3. Markus G Rudolph, Robyn L Stanfield, and Ian A Wilson. How tcrs bind mhcs, peptides, and coreceptors. Annu. Rev. Immunol., 24(1):419–466, 2006.

4. K Christopher Garcia and Erin J Adams. How the t cell receptor sees antigen—a structural view. Cell, 122(3):333–336, 2005.

5. Michael E Birnbaum, Juan L Mendoza, Dhruv K Sethi, Shen Dong, Jacob Glanville, Jessica Dobbins, Engin Özkan, Mark M Davis, Kai W Wucherpfennig, and K Christopher Garcia. Deconstructing the peptide-mhc specificity of t cell recognition. Cell, 157(5):1073–1087, 2014.

6. Michael A Morse, William R Gwin III, and Duane A Mitchell. Vaccine therapies for cancer: then and now. Targeted oncology, 16(2):121–152, 2021.

7. Cassian Yee. Adoptive t cell therapy: addressing challenges in cancer immunotherapy. Journal of Translational Medicine, 3(1):1–8, 2005.

8. Maria Harkiolaki, Samantha L Holmes, Pia Svendsen, Jon W Gregersen, Lise T Jensen, Roisin McMahon, Manuel A Friese, Gijs Van Boxel, Ruth Etzensperger, John S Tzartos, et al. T cell-mediated autoimmune disease due to low-affinity crossreactivity to common microbial peptides. Immunity, 30(3):348–357, 2009.

9. Jonathan D Buhrman, Kimberly R Jordan, Daniel J Munson, Brandon L Moore, John W Kappler, and Jill E Slansky. Improving antigenic peptide vaccines for cancer immunotherapy using a dominant tumor-specific t cell receptor. Journal of Biological Chemistry, 288(46): 33213–33225, 2013.

10. Geir Åge Løset, Gøril Berntzen, Terje Frigstad, Sylvie Pollmann, Kristin S Gunnarsen, and Inger Sandlie. Phage display engineered t cell receptors as tools for the study of tumor peptide–mhc interactions. Frontiers in oncology, 4:378, 2015.

11. Marvin H Gee, Arnold Han, Shane M Lofgren, John F Beausang, Juan L Mendoza, Michael E Birnbaum, Michael T Bethune, Suzanne Fischer, Xinbo Yang, Raquel Gomez-Eerland, et al. Antigen identification for orphan t cell receptors expressed on tumor-infiltrating lymphocytes. Cell, 172(3):549–563, 2018.

12. Ido Springer, Hanan Besser, Nili Tickotsky-Moskovitz, Shirit Dvorkin, and Yoram Louzoun. Prediction of specific tcr-peptide binding from large dictionaries of tcr-peptide pairs. Frontiers in immunology, page 1803, 2020.

13. David N Garboczi, Partho Ghosh, Ursula Utz, Qing R Fan, William E Biddison, and Don C Wiley. Structure of the complex between human t-cell receptor, viral peptide and hla-a2. Nature, 384(6605):134–141, 1996.

14. David H Margulies, Daniel Plaksin, SN Khilko, and Marie T Jelonek. Studying interactions involving the t-cell antigen receptor by surface plasmon resonance. Current opinion in immunology, 8(2):262–270, 1996.

15. Eric T Boder, Jerome R Bill, Andrew W Nields, Philippa C Marrack, and John W Kappler. Yeast surface display of a noncovalent mhc class ii heterodimer complexed with antigenic peptide. Biotechnology and bioengineering, 92(4):485–491, 2005.

16. Wei Jiang and Eric T Boder. High-throughput engineering and analysis of peptide binding to class ii mhc. Proceedings of the National Academy of Sciences, 107(30):13258–13263, 2010.

17. Scott E Starwalt, Emma L Masteller, Jeffrey A Bluestone, and David M Kranz. Directed evolution of a single-chain class ii mhc product by yeast display. Protein engineering, 16(2): 147–156, 2003.

18. Xingang Peng, Yipin Lei, Peiyuan Feng, Lemei Jia, Jianzhu Ma, Dan Zhao, and Jianyang Zeng. Characterizing the interaction conformation between t-cell receptors and epitopes with deep learning. Nature Machine Intelligence, pages 1–13, 2023.

19. Dan Hudson, Ricardo A Fernandes, Mark Basham, Graham Ogg, and Hashem Koohy. Can we predict t cell specificity with digital biology and machine learning? Nature Reviews Immunology, 23(8):511–521, 2023.

20. Chloe H Lee, Jaesung Huh, Paul R Buckley, Myeongjun Jang, Mariana Pereira Pinho, Ricardo A Fernandes, Agne Antanaviciute, Alison Simmons, and Hashem Koohy. A robust deep learning workflow to predict cd8+ t-cell epitopes. Genome Medicine, 15(1):70, 2023.

21. Zahra S Ghoreyshi and Jason T George. Quantitative approaches for decoding the specificity of the human t cell repertoire. Frontiers in Immunology, 14:1228873, 2023.

22. Wen Zhang, Peter G Hawkins, Jing He, Namita T Gupta, Jinrui Liu, Gabrielle Choonoo, Se W Jeong, Calvin R Chen, Ankur Dhanik, Myles Dillon, et al. A framework for highly multiplexed dextramer mapping and prediction of t cell receptor sequences to antigen specificity. Science Advances, 7(20):eabf5835, 2021.

23. Vanessa Isabell Jurtz, Leon Eyrich Jessen, Amalie Kai Bentzen, Martin Closter Jespersen, Swapnil Mahajan, Randi Vita, Kamilla Kjærgaard Jensen, Paolo Marcatili, Sine Reker Hadrup, Bjoern Peters, et al. Nettcr: sequence-based prediction of tcr binding to peptidemhc complexes using convolutional neural networks. BioRxiv, page 433706, 2018.

24. Chloe H Lee, Mariolina Salio, Giorgio Napolitani, Graham Ogg, Alison Simmons, and Hashem Koohy. Predicting cross-reactivity and antigen specificity of t cell receptors. Frontiers in Immunology, 11:565096, 2020.

25. Galina Petrova, Andrea Ferrante, and Jack Gorski. Cross-reactivity of t cells and its role in the immune system. Critical Reviews™ in Immunology, 32(4), 2012.

26. Dinler A Antunes, Maurício M Rigo, Martiela V Freitas, Marcus FA Mendes, Marialva Sinigaglia, Gregory Lizée, Lydia E Kavraki, Liisa K Selin, Markus Cornberg, and Gustavo F Vieira. Interpreting t-cell cross-reactivity through structure: implications for tcr-based cancer immunotherapy. Frontiers in immunology, 8:1210, 2017.

27. Xingcheng Lin, Jason T George, Nicholas P Schafer, Kevin Ng Chau, Michael E Birnbaum, Cecilia Clementi, José N Onuchic, and Herbert Levine. Rapid assessment of t-cell receptor specificity of the immune repertoire. Nature Computational Science, 1(5):362–373, 2021.

28. Ailun Wang, Xingcheng Lin, Kevin Ng Chau, José N Onuchic, Herbert Levine, and Jason T George. Racer-m leverages structural features for sparse t cell specificity prediction. Science Advances, 10(20):eadl0161, 2024.

29. Zahra S Ghoreyshi, Hamid Teimouri, Anatoly T Kolomeisky, and Jason T George. Integration of kinetic data into affinity-driven models for improved t cell-antigen specificity prediction. bioRxiv, pages 2024–06, 2024.

30. Barthelemy Meynard-Piganeau, Christoph Feinauer, Martin Weigt, Aleksandra M Walczak, and Thierry Mora. Tulip: A transformer-based unsupervised language model for interacting peptides and t cell receptors that generalizes to unseen epitopes. Proceedings of the National Academy of Sciences, 121(24):e2316401121, 2024.

31. Pradyot Dash, Andrew J Fiore-Gartland, Tomer Hertz, George C Wang, Shalini Sharma, Aisha Souquette, Jeremy Chase Crawford, E Bridie Clemens, Thi HO Nguyen, Katherine Kedzierska, et al. Quantifiable predictive features define epitope-specific t cell receptor repertoires. Nature, 547(7661):89–93, 2017.

32. Jacob Glanville, Huang Huang, Allison Nau, Olivia Hatton, Lisa E Wagar, Florian Rubelt, Xuhuai Ji, Arnold Han, Sheri M Krams, Christina Pettus, et al. Identifying specificity groups in the t cell receptor repertoire. Nature, 547(7661):94–98, 2017.

33. Alessandro Montemurro, Viktoria Schuster, Helle Rus Povlsen, Amalie Kai Bentzen, Vanessa Jurtz, William D Chronister, Austin Crinklaw, Sine R Hadrup, Ole Winther, Bjoern Peters, et al. Nettcr-2.0 enables accurate prediction of tcr-peptide binding by using paired tcrα and β sequence data. Communications biology, 4(1):1060, 2021.

34. Evan W Newell, Lauren K Ely, Andrew C Kruse, Philip A Reay, Stephanie N Rodriguez, Aaron E Lin, Michael S Kuhns, K Christopher Garcia, and Mark M Davis. Structural basis of specificity and cross-reactivity in t cell receptors specific for cytochrome c–i-ek. The Journal of Immunology, 186(10):5823–5832, 2011.

35. Haibo He and Edwardo A Garcia. Learning from imbalanced data. IEEE Transactions on knowledge and data engineering, 21(9):1263–1284, 2009.

36. Dong-Sheng Cao, Qing-Song Xu, and Yi-Zeng Liang. propy: a tool to generate various modes of chou’s pseaac. Bioinformatics, 29(7):960–962, 2013.

37. Hamid Teimouri, Angela Medvedeva, and Anatoly B Kolomeisky. Bacteria-specific feature selection for enhanced antimicrobial peptide activity predictions using machine-learning methods. Journal of Chemical Information and Modeling, 63(6):1723–1733, 2023.

38. Ernest Y Lee, Michelle W Lee, Benjamin M Fulan, Andrew L Ferguson, and Gerard CL Wong. What can machine learning do for antimicrobial peptides, and what can antimicrobial peptides do for machine learning? Interface focus, 7(6):20160153, 2017.

39. Hamid Teimouri, Angela Medvedeva, and Anatoly B Kolomeisky. Unraveling the role of physicochemical differences in predicting protein–protein interactions. The Journal of Chemical Physics, 161(4), 2024.

40. William B Cavnar, John M Trenkle, et al. N-gram-based text categorization. In Proceedings of SDAIR-94, 3rd annual symposium on document analysis and information retrieval, volume 161175, page 14. Ann Arbor, Michigan, 1994.

41. Zhiqiang Zeng, Hua Shi, Yun Wu, and Zhiling Hong. Survey of natural language processing techniques in bioinformatics. Computational and mathematical methods in medicine, 2015 (1):674296, 2015.

42. Hui Ding, Hao Lin, Wei Chen, Zi-Qiang Li, Feng-Biao Guo, Jian Huang, and Nini Rao. Prediction of protein structural classes based on feature selection technique. Interdisciplinary Sciences: Computational Life Sciences, 6:235–240, 2014.

43. SM Ashiqul Islam, Benjamin J Heil, Christopher Michel Kearney, and Erich J Baker. Protein classification using modified n-grams and skip-grams. Bioinformatics, 34(9):1481–1487, 2018.

44. Robert Tibshirani. Regression shrinkage and selection via the lasso. Journal of the Royal Statistical Society Series B: Statistical Methodology, 58(1):267–288, 1996.

45. Vincent Detours, Ramit Mehr, and Alan S Perelson. A quantitative theory of affinity-driven t cell repertoire selection. Journal of theoretical biology, 200(4):389–403, 1999.

46. Andrej Košmrlj, Abhishek K Jha, Eric S Huseby, Mehran Kardar, and Arup K Chakraborty. How the thymus designs antigen-specific and self-tolerant t cell receptor sequences. Proceedings of the National Academy of Sciences, 105(43):16671–16676, 2008.

47. Jason T George, David A Kessler, and Herbert Levine. Effects of thymic selection on t cell recognition of foreign and tumor antigenic peptides. Proceedings of the National Academy of Sciences, 114(38):E7875–E7881, 2017.

48. Kevin Ng Chau, Jason T George, José N Onuchic, Xingcheng Lin, and Herbert Levine. Contact map dependence of a t-cell receptor binding repertoire. Physical Review E, 106(1): 014406, 2022.

49. Philip Bradley. Structure-based prediction of t cell receptor: peptide-mhc interactions. eLife, 12:e82813, 2023.

50. Michael E Birnbaum, Juan L Mendoza, Dhruv K Sethi, Shen Dong, Jacob Glanville, Jessica Dobbins, Engin Özkan, Mark M Davis, Kai W Wucherpfennig, and K Christopher Garcia. Deconstructing the peptide-mhc specificity of t cell recognition. Cell, 157(5):1073–1087, 2014.

