## Supplementary Information for "Feature Selection Enhances Peptide Binding Predictions for TCR-Specific Interactions"

### Supplementary Figures

In this supporting information, we present additional figures that are discussed in the main text.

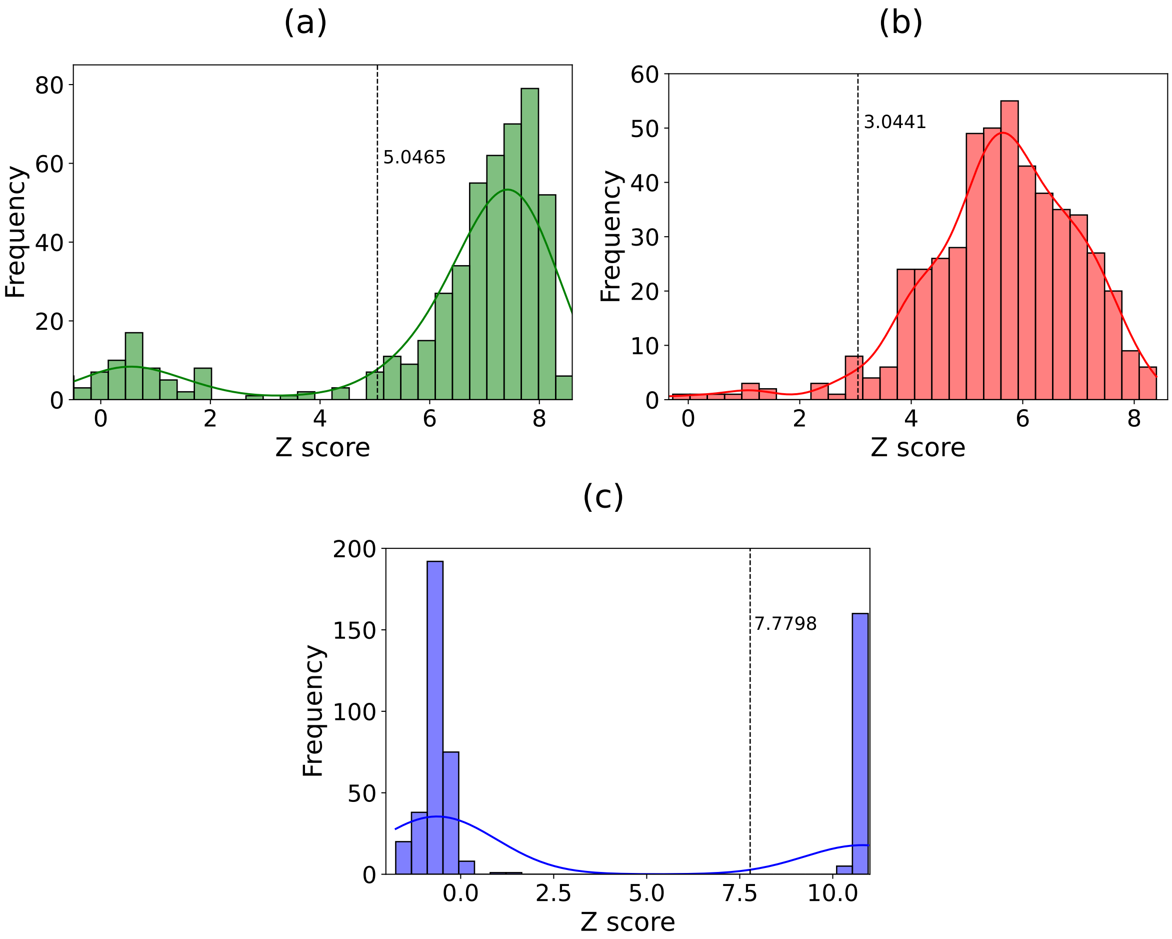


Figure S1: Calculation of threshold value suing RACER model for peptide libraries associate with (a) 2B4 TCR , (b) 226 TCR , (c) 5cc7 TCR.

### Histograms for Calculating Thresholds

Corresponding histograms for calculating post selection abundance threshold, which separates strong binders from weak binders, are presented in Fig. S1.

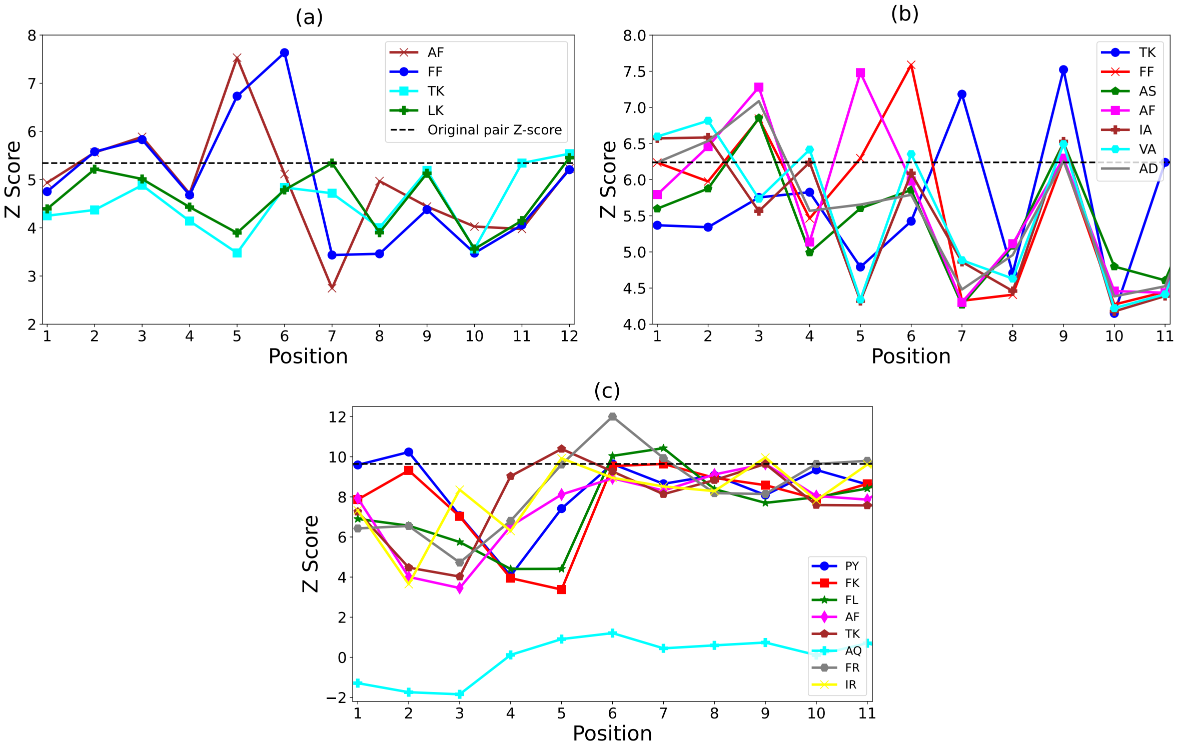


Figure S2: RACER model predictions for the selected dipeptide compositions for (a) 2B4, (b) 226, (c) 5cc7.

### Dipeptides from RACER model

The corresponding position curves, calculated by the RACER model for dipeptide feature selected by LASSO, are presented in Fig. S2.

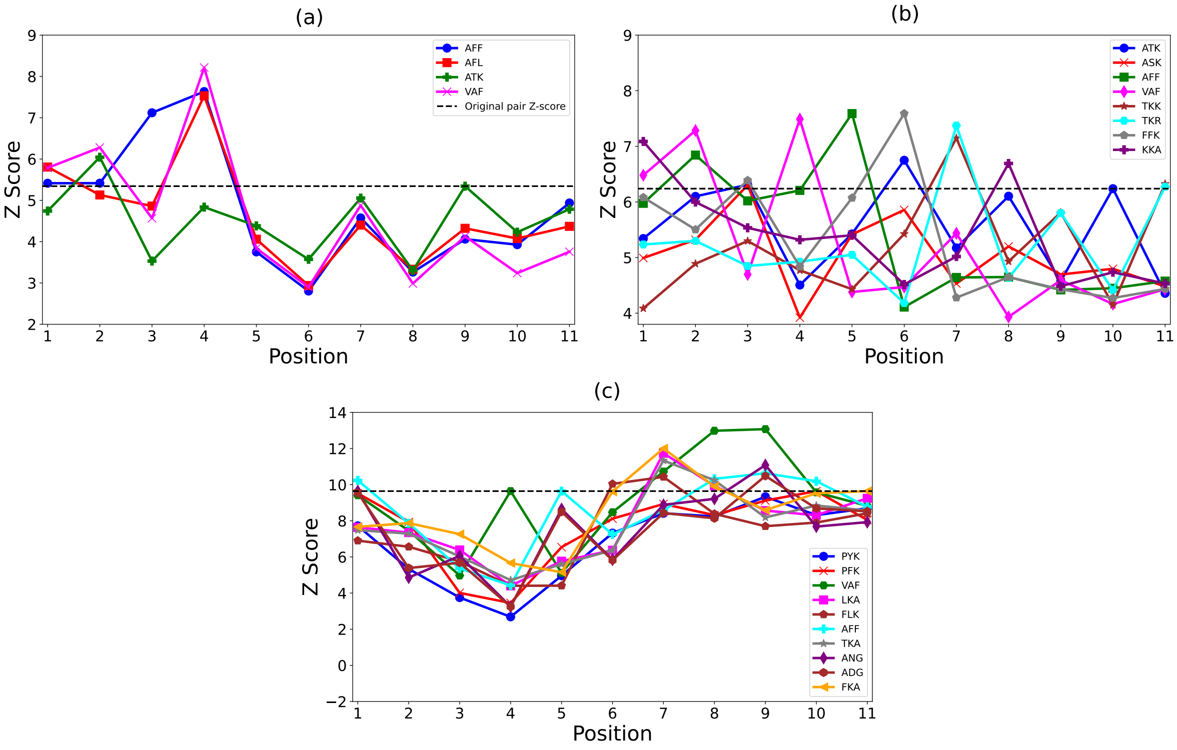


Figure S3: RACER model predictions for the selected tripeptide compositions for (a) 2B4, (b) 226, (c) 5cc7.

### Tripeptide from RACER model

Fig. S3 displays the position curves for tripeptide features selected by LASSO, calculated using the RACER model.

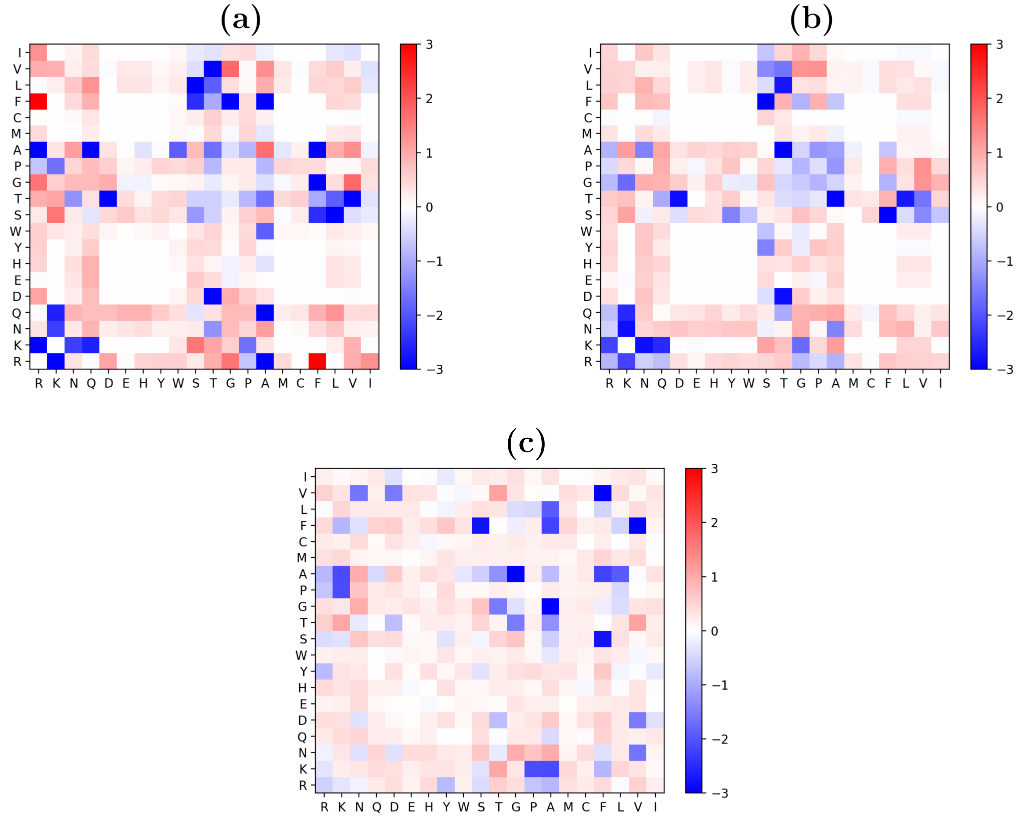


Figure S4: The residue-based interaction strength, represented by the energy matrix, determined by RACER with a maximum distance of $r_{\mathrm{max}} = 8.5$Åfor TCR (a) 2B4, (b) 226, and (c) 5cc7.

### Analysis of TCR-Peptide contact maps and energy matrices

In this study, we present RACER-driven energy matrices and corresponding contact maps for various TCR-peptide complexes, all generated using a maximum distance cutoff of 8.5Å (Fig. S4 and S5).

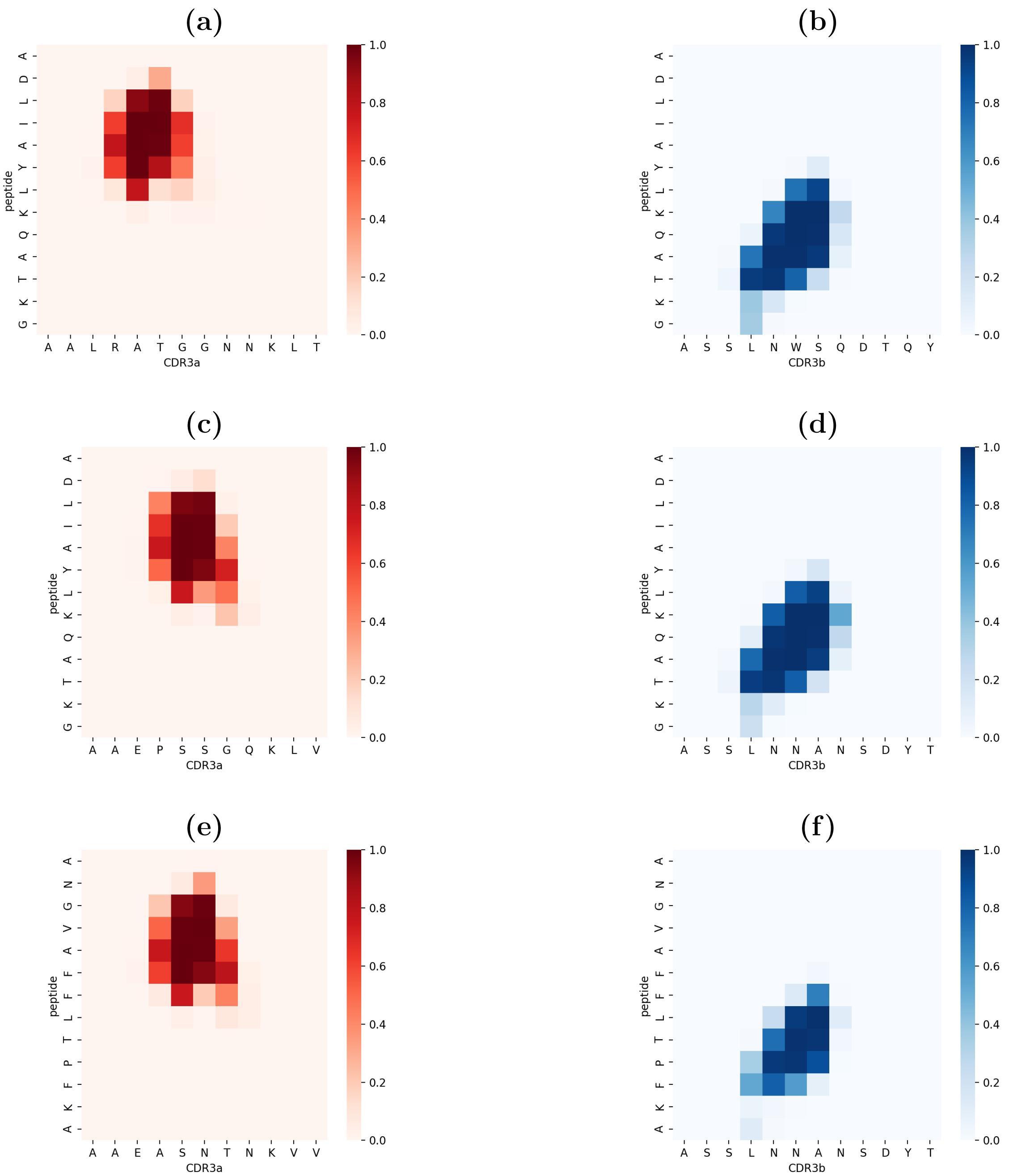


Figure S5: Contact maps illustrating the peptide-TCR interactions for different TCR-peptide complexes, generated using a maximum distance ($r_{\max} = 8.5$Å): **(a)** 2B4 CDR3$\alpha$, **(b)** 2B4 CDR3$\beta$, **(c)** 226 CDR3$\alpha$, **(d)** 226 CDR3$\beta$, **(e)** 5cc7 CDR3$\alpha$, and **(f)** 5cc7 CDR3$\beta$.

Table S1: Description of selected *propy* features related to Figs 3(a) and Fig 4(a) in the main text.

| **Feature Acronym** | **Feature Description** |
|---|---|
| PolarityD2075 | The fraction of the entire sequence, where 75% of the residues of group 2 (polarity values $8.0 - 9.2$) are contained. |
| GearyAuto_Hydrophobicity5 | Geary's autocorrelation function of hydrophobicity for amino acids that are 5 positions apart. |
| QSOSW12 | Quasi-sequence order |
| QSOgrant5 | Quasi-order-coupling number |
| NormalizedVDWVC2 | Global percent composition of residues with normalized van der Waals volume in range $2.95 - 94.0$. |
